## supplementary tables for "Toward perfect reads: short reads correction via mapping on compacted de Bruijn graphs"

*Bioinformatics*
doi.10.1093/bioinformatics/xxxxxx
Advance Access Publication Date: Day Month Year
Manuscript Category

OXFORD

### Supplementary materials

**Antoine Limasset[1], Jean-François Flot[1,2,*], and Pierre Peterlongo[3,*]**

[1]Université libre de Bruxelles (ULB), Evolutionary Biology & Ecology, B-1050 Bruxelles, Belgium
[2]Interuniversity Institute of Bioinformatics in Brussels – (IB)$^2$, Brussels, Belgium
[3]Univ Rennes, Inria, CNRS, IRISA, F-35000 Rennes, France

*Co-last authors with equal contributions.

Associate Editor: D

Received on D; revised on D; accepted on D

#### Abstract

Sections 1 to 4 provide clarifications and additional results to those shown in Section 3.2 (longer reads, lower coverage, lower error rate, distinct $k$ values, diploid simulations). For all results except those presented in Table 2 we used the default $k$ value of each tool. All simulations (except in Section 3) are directly made from reference genomes and do not contain diploid variations.
Section 5 provides additional information on the tipping and unitig-filtering strategies.

For all presented results, the sensitivity is given by $\frac{TP}{TP+FN}$ and the specificity by $\frac{TN}{TN+FP}$.

#### 1 Results on simulated *C. elegans* data

In this section we provide additional results obtained on *C. elegans* for various read lengths and coverage depths (Table 1). We also performed tests using a 0.5% error rate and obtained similar results (data not shown). Additionally, we provide results (Table 2) obtained while using a high $k$ value.

#### 2 Results on simulated human chromosome 1 data

In this section we provide additional results obtained on the human chromosome 1 for various read lengths and coverage depths (Table 3). We also performed tests using a 0.5% error rate and obtained similar results (data not shown).

| Corrector | Sensitivity | Specificity | Correction ratio | % Erroneous reads |
|---|---|---|---|---|
| 150-bp reads at 100X coverage | | | | |
| Bcool | **99.595** | **99.999** | **190.793** | **0.303** |
| BFC | 94.854 | 99.997 | 18.53 | 3.634 |
| Bloocoo | 96.852 | 99.993 | 26.14 | 3.753 |
| Lighter | 97.352 | 99.995 | 31.70 | 2.857 |
| Musket | 98.922 | 99.997 | 73.89 | 1.466 |
| 150-bp reads at 50X coverage | | | | |
| Bcool | **99.467** | **99.999** | **151.35** | **0.421** |
| BFC | 95.857 | 99.998 | 22.98 | 2.789 |
| Bloocoo | 97.090 | 99.994 | 28.14 | 3.5 |
| Lighter | 98.149 | 99.996 | 43.97 | 1.996 |
| Musket | 98.822 | 99.997 | 68.68 | 1.569 |
| 250-bp reads at 100X coverage | | | | |
| Bcool | **99.537** | **99.999** | **183.53** | **0.458** |
| Bloocoo | 97.376 | 99.993 | 30.28 | 4.893 |
| BFC | 93.327 | 99.998 | 14.58 | 5.667 |
| Lighter | 97.346 | 99.994 | 31.16 | 4.225 |
| Musket | 99.142 | 99.998 | 90.33 | 1.867 |
| 250-bp reads at 50X coverage | | | | |
| Bcool | **99.498** | **99.999** | **162.98** | **0.516** |
| Bloocoo | 97.634 | 99.994 | 33.34 | 4.509 |
| BFC | 94.541 | 99.998 | 17.82 | 4.431 |
| Lighter | 98.008 | 99.995 | 40.28 | 3.203 |
| Musket | 99.071 | 99.998 | 84.70 | 1.963 |

Table 1. Correction metrics of various correctors applied on C. elegans reads simulated with a 1% error rate

#### 3 Results on simulated human chromosome 1 diploid data

Two vcf files were retrieved from the “1000 genome project” (phase 1 release), corresponding to the human chromosome 1 of two individuals: HG00096 and HG00100. We then generated the genome sequences for the two diploids, i.e. two sequences per individual, by placing the substitutions listed in the vcf files onto the human reference sequence (GRCh37/hg19 reference assembly version). A total of 316,502 positions were mutated, with 131,263 positions mutated at the same time in both individuals (representing an average 0.5 SNP per Kb in each individual). 29,038 SNPs (9 %) were homozygous in both individuals (homozygous-homozygous),

© The Author 2018. Published by Oxford University Press. All rights reserved. For permissions, please

| Corrector | Sensitivity | Specificity | Correction ratio | % Erroneous reads |
|---|---|---|---|---|
| $k = 63$ | | | | |
| Bcool | **99.395** | **99.998** | **129.504** | **0.519** |
| Bloocoo | 82.933 | 99.992 | 5.590 | 14.210 |
| Lighter | 96.598 | **99.998** | 27.493 | 1.5276 |
| $k = 95$ | | | | |
| Bcool | **99.590** | **99.999** | **178.904** | **0.321** |
| Bloocoo | 63.5537 | 99.997 | 2.722 | 20.116 |
| Lighter | 61.240 | 99.995 | 2.547 | 25.458 |

Table 2. Simulated C. elegans 150-bp reads with 1% error rate and 100X coverage/ The Musket run was not able to complete and BFC yielded a correction ratio $< 1$ - both are therefore not reported here.

| Corrector | Sensitivity | Specificity | Correction ratio | % Erroneous reads |
|---|---|---|---|---|
| 150-bp reads at 100X coverage | | | | |
| Bcool | **99.017** | **99.999** | **95.40** | **0.745** |
| BFC | 86.225 | 99.991 | 6.82 | 10.519 |
| Bloocoo | 92.573 | 99.965 | 9.15 | 9.03 |
| Lighter | 91.269 | 99.975 | 8.96 | 8.709 |
| Musket | 93.052 | 99.982 | 11.4 | 8.307 |
| 150-bp reads at 50X coverage | | | | |
| Bcool | **98.193** | **99.998** | **48.83** | **1.505** |
| BFC | 87.397 | 99.991 | 7.434 | 9.543 |
| Bloocoo | 93.053 | 99.962 | 9.38 | 8.641 |
| Lighter | 91.915 | 99.976 | 9.550 | 8.123 |
| Musket | 92.876 | 99.982 | 11.19 | 8.458 |
| 250-bp reads at 100X coverage | | | | |
| Bcool | **99.392** | **100** | **153.73** | **0.577** |
| Bloocoo | 93.291 | 99.964 | 9.76 | 12.308 |
| Lighter | 90.336 | 99.977 | 8.35 | 13.616 |
| Musket | 93.816 | 99.982 | 12.6 | 10.708 |
| BFC | 82.744 | 99.994 | 5.6 | 15.546 |
| 250-bp reads at 50X coverage | | | | |
| Bcool | **98.855** | **99.999** | **77.376** | **1.214** |
| Bloocoo | 93.774 | 99.962 | 10.038 | 11.798 |
| Lighter | 91.717 | 99.977 | 9.42 | 11.855 |
| BFC | 84.008 | 99.994 | 6.05 | 14.170 |
| Musket | 93.666 | 99.982 | 12.372 | 10.864 |

Table 3. Correction metrics of various correctors applied on human chromosome 1 reads simulated with a 1% error rate

| Corrector | Sensitivity | Specificity | Correction ratio | % Erroneous reads |
|---|---|---|---|---|
| 250-bp reads at 100X coverage | | | | |
| Bcool | **99.506** | **99.998** | **152.60** | **0.685** |
| Bloocoo | 94.045 | 99.966 | 10.71 | 11.415 |
| BFC | 83.996 | 99.994 | 6.04 | 14.256 |
| Musket | 94.314 | 99.982 | 13.461 | 10.298 |
| Lighet | 91.309 | 99.977 | 9.143 | 12.648 |
| 150-bp reads at 100X coverage | | | | |
| Bcool | **99.256** | **99.998** | **100.40** | **0.720** |
| Bloocoo | 93.330 | 99.966 | 9.98 | 8.281 |
| BFC | 87.438 | 99.992 | 7.472 | 9.474 |
| Musket | 93.548 | 99.982 | 12.11 | 7.907 |
| Lighter | 91.414 | 99.977 | 9.176 | 8.667 |

Table 4. Correction metrics of various correctors applied on diploid human chromosome1 reads simulated with 1% error rate

| Corrector | Sensitivity | Specificity | Correction ratio | % Erroneous reads |
|---|---|---|---|---|
| 150-bp reads at 100X coverage | | | | |
| Bcool | **97.735** | **99.996** | **37.73** | **1.946** |
| Bloocoo | 88.901 | 99.956 | 6.48 | 12.427 |
| Lighter | 85.565 | 99.971 | 5.78 | 13.639 |
| 150-bp reads at 50X coverage | | | | |
| Bcool | **96.495** | **99.996** | **25.66** | **2.916** |
| Bloocoo | 89.591 | 99.953 | 6.63 | 11.962 |
| Lighter | 87.414 | 99.97 | 6.42 | 11.954 |
| 250-bp reads at 100X coverage | | | | |
| Bcool | **98.415** | **99.998** | **56.80** | **1.718** |
| Bloocoo | 89.649 | 99.956 | 6.789 | 16.810 |
| Lighter | 84.969 | 99.973 | 5.65 | 19.639 |
| 250-bp reads at 50X coverage | | | | |
| Bcool | **97.621** | **99.996** | **35.87** | **2.668** |
| Bloocoo | 90.366 | 99.953 | 6.98 | 16.200 |
| Lighter | 87.109 | 99.971 | 6.35 | 16.964 |

Table 5. Correction metrics of various correctors applied to reads simulated from the complete human genome with a 1% error rate

218,556 (69 %) were heterozygous in only one individual (homozygous-heterozygous) and the remaining 68,908 (22 %) were heterozygous in both individuals. We then simulated a 100X coverage sequencing with a 1% error rate from this pair of diploid genomes. Results are presented Table 4.

#### 4 Results on simulated whole human genome data

In this section we provide additional results obtained on the whole human genome for various read lengths and sequencing depths (Table 5). We also performed tests using a 0.5% error rate and obtained similar results (data not shown).

#### 5 DBG construction strategies

In this section we evaluate diverse DBG construction strategies. We performed tests on a simulated *C. elegans* 50X data set of 150-bp reads with a 1% error rate. Results are presented in Table 6. We show only graph-cleaning results obtained with low $k$-mer abundance thresholds (2 and 3) as higher values would not make sense for our approach. Results using higher $k$-mer abundance thresholds are showed here for KAF only, as it corresponds to a classical $k$-mer spectrum approach.

| $k$-mer abundance threshold | KAF | KAF+TIP | KAF+ UAF | KAF+TIP+UAF |
|---|---|---|---|---|
| k=31 | | | | |
| 2 | 54,075,339 / 36 | 19,429,473 / 550 | 858,726 / 46 | 642,396 / 560 |
| 3 | 5,628,920 / 55 | 857,469 / 57 | 676,110 / 623 | 676,110 / 623 |
| 4 | 1,968,288 / 75 | | | |
| 5 | 1,115,357 / 85 | | | |
| 10 | 216,094 / 1,175 | | | |
| k=63 | | | | |
| 2 | 31,902,775 / 347 | 1,586,655 / 2,070 | 127,975 / 1,409 | 78,639 / 3,963 |
| 3 | 1,837,789 / 2,507 | 176,693 / 4,612 | 136,972 / 3,295 | 76,407 / 4,653 |
| 4 | 482,095 / 13,145 | | | |
| 5 | 217,508 / 54,225 | | | |
| 10 | 17,508 / 5,083,037 | | | |

Table 6. Evaluation of different DBG construction strategies proposed by Bcool. Each result presents two values ($v_1$/$v_2$). Value $v_1$ is the number of erroneous $k$-mers present in the DBG and $v_2$ is the number of genomic $k$-mers missing in the DBG. In this experiment the unitig filtering threshold was set to five. The different rows represent the efficiency of the different strategies tested: $k$-mer abundance filter alone (KAF), tip removal after $k$-mer filter (KAF+TIP), unitig abundance filtering after $k$-mer filtering (KAF+UAF), and the combination of the three strategies (KAF+TIP+UAF).
